# Pilot Suppression trial of *Aedes albopictus* mosquitoes through an Integrated Vector Management strategy including the Sterile Insect Technique in Mauritius

**DOI:** 10.1101/2020.09.06.284968

**Authors:** Diana P. Iyaloo, Jeremy Bouyer, Sunita Facknath, Ambicadutt Bheecarry

## Abstract

It is often difficult to control the vector mosquito *Aedes albopictus* using conventional chemical control methods alone at an operational level mainly because of (1) the ability of the species to lay eggs in a variety of places which are often difficult to detect or access by larviciding operators, (2) the inherent tendency of adults to live and feed outdoor which makes them unlikely targets of Insecticide Residual Spraying and (3) the development of resistance to insecticides by the species. It is therefore necessary for countries to investigate alternative control methods (such as the Sterile Insect Technique (SIT)) that can be integrated in their national vector control programme in order to address those limitations.

In this field trial, mass-produced, radio-sterilized *Ae. albopictus* males could successfully compete with wild males in a small village in Mauritius. Our study also demonstrated that within specific eco-climatic conditions, SIT can be used as a suppression tool against *Ae. albopictus* and, unlike numerous chemical control methods, effectively maintain the suppression level when the latter is found at low densities. Finally, the need for mosquito SIT programmes to develop contingency plans against increasingly frequent extreme weather occurrences was also highlighted.

## Introduction

The control of *Aedes albopictus* is a high priority for the Mauritian Government since the species is the only vector responsible in the transmission of Chikungunya (a severe outbreak in 2005-2006) and Dengue (outbreaks in 2009, 2014, 2015, 2019 and 2020) in the country^1,2,3,4,5,6^. After the eradication of *Aedes aegypti* in the 1950s, the species has since then never detected on the island during routine mosquito surveys carried out by the Vector Biology and Control Division (a national department specialized in mosquito surveillance). Despite numerous measures currently undertaken by the Mauritian Health authorities to control the species (including regular larviciding of mosquito breeding sites, bi-annual spraying of ports of entries with a pyrethroid-based insecticide and fogging of patient’s locality following confirmed or suspected cases of vector-borne diseases), it is still difficult to control *Ae. albopictus* since the species breeds in a wide variety of artificial and natural water-collecting containers which are often difficult to detect or access by larviciding operators. Furthermore, besides the numerous limitations linked to an effective field application of chemical insecticides, the latter may also cause adverse effects at different levels of the ecosystem ranging from bioaccumulation, lethal effects on non-target fauna and resistance build-up in the mosquito vector^7^. Conscious that the Integrated Vector Management strategy would need to be reinforced for a sustainable control of *Ae. albopictus* in the country, investigation of the Sterile Insect Technique (SIT) as an additional control tool is encouraged by the Mauritian Authority in view of its potential inclusion in the national mosquito control programme.

The SIT is a method of pest control using area-wide inundative releases of sterile insects to reduce reproduction in a field population of the same species^8^. Although the principle of SIT is quite straight-forward, many challenges are linked to its effective application in the field^9^. For instance, the release of 180,000 sterile males of *Culex tarsalis* over a period of 2.5 months in California failed to reduce its wild population due to the low competitiveness and dispersal of the sterile males^10^. Sterile male release of *Culex tritaeniorhynchus* in Pakistan (167,000 males over a 2-week period) also did not reduce its wild population due to the inability of the sterile males to compete for wild females and because of immigration of fertile females into the test sites^11^. Similarly, in El Salvador, unexpected immigration was believed to have mitigated the suppression programme of *Anopheles albimanus* where more than 100 million sterile males were released over 3 years^12^). The poor competitiveness of sterile males of *Anopheles gambiae* released in Burkina Faso (240,000 sterile males) over 9 weeks was also the cause of the failure of the suppression programme^13^. On the other hand, in two consecutive release programmes, sterile males of *Anopheles quadrimaculatus* in Florida could not induce sterility in the target population due to behavioral differences as a result of the rearing process^14, 15^.

The aim of this study was hence to investigate whether the SIT could be used to control the wild population of *Ae. albopictus* in Panchvati (PAN), a small village in the northeast of Mauritius.

## Results

### Effects of climate

Ovitrap productivity followed a seasonal trend in the release site PAN, within a radius of 150 m around PAN (henceforth referred to as ‘PAN_Out’) and in the control site Pointe des Lascars (henceforth referred to as ‘PDL’). Ovitrap productivity remained relatively higher in the three sites during the hot and humid summer season (spanning from November to April) as compared to the winter season (May to October) (Fig. 1b and d). In PAN and PDL, a highly significant positive correlation was noted between weekly mean of ovitrap productivity and temperature (*P* < 0.0001) but not with cumulative rainfall (P > 0.05) while in PAN_Out, ovitrap productivity was significantly positively correlated with both temperature (*P* < 0.0001) and rainfall (*P* = 0.0037) (Table 1).

**Figure 1.**
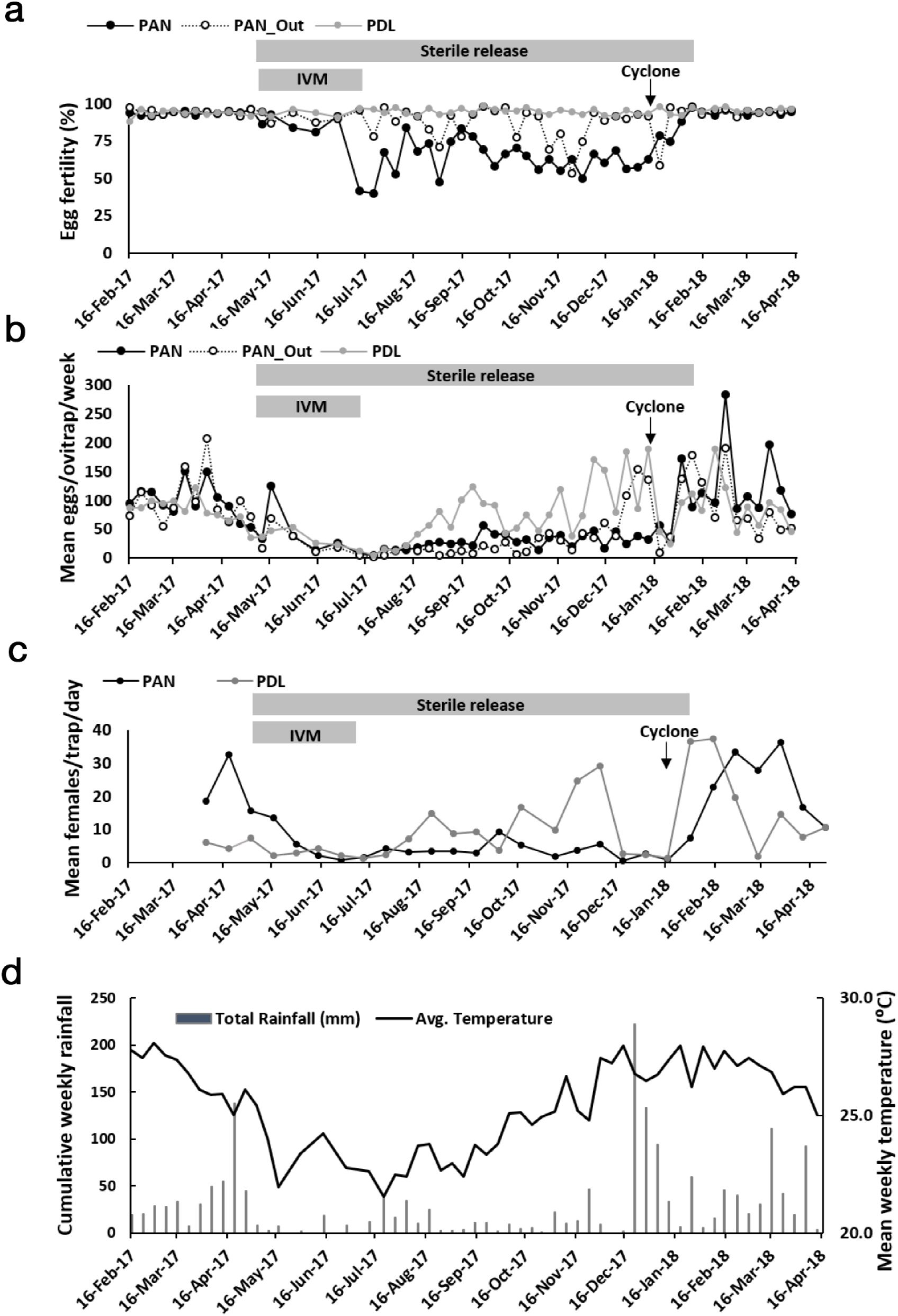
**(a)** Weekly mean of fertility rate and **(b)** density of *Ae. albopictus* eggs in ovitraps, **(c)** daily capture rate of *Ae. albopictus* females from BGS traps in PDL, PAN and PAN_Out and **(d)** weekly cumulative rainfall and mean air temperature for the region

**Table 1.** Spearman’s rank correlation coefficient (rs) between weather data (weekly mean of temperature and cumulative rainfall), ovitrap productivity (eggs/ovitrap/week) and female *Ae. albopictus* capture rate from BGS traps in PAN, PAN_Out and PDL from April 2017 to April 2018.

| Sites | Weather<br>variable | Ovitrap |  |
| --- | --- | --- | --- |
| | | productivity<br>$r_s$ | Female capture<br>$r_s$ |
| PDL | Temperature | 0.57, (n = 57, $P <$ | 0.24, (n = 27, $P =$ |
|  |  | 0.0001) | 0.219) |
| PAN | | 0.58, (n = 57, $P <$ | 0.17, (n = 27, $P =$ |
|  |  | 0.0001) | 0.40) |
| PAN_Out | | 0.68, (n = 57, $P <$ | Not available |
|  |  | 0.0001) |  |
| PDL | Rainfall | 0.07, (n = 57, $P =$ | -0.07, (n = 27, $P =$ |
|  |  | 0.591) | 0.745) |
| PAN | | 0.20, (n = 57, $P =$ | 0.24, (n = 27, $P =$ |
|  |  | 0.145) | 0.238) |
| PAN_Out | | 0.38, (n = 57, $P =$ | Not available |
|  |  | 0.0037) |  |

### Impact of the IVM strategy

From mid-May to mid-July 2017, the implementation of an IVM strategy coupled with the beginning of the unfavorable dry winter season (Fig 1d), led to an approximate 3-fold reduction in ovitrap productivity in PAN and PDL (Fig. 1b, Table 2). An important reduction in *Ae. albopictus* female collection rate by BG Sentinel™ traps (henceforth referred as ‘BGS traps’) was also noted (Fig. 1c), dropping from a mean (95 % CI) of 22 (12-33) and 6 (4-8) females/BGS trap/day in PAN and PDL respectively prior to the start of the IVM strategy to 5 (1-8) and 2 (2-3) females/BGS trap/day when IVM was implemented.

**Table 2.**
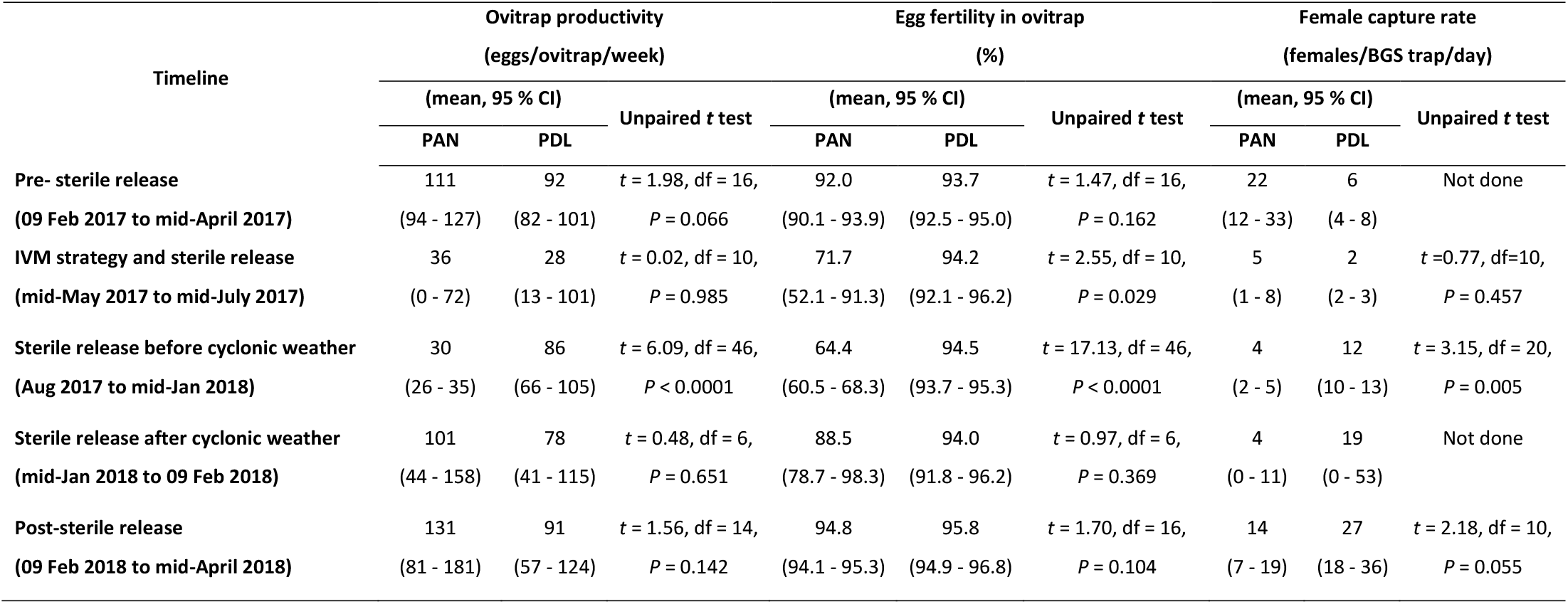
Mean weekly ovitrap productivity, mean weekly egg fertility and mean daily adult female capture rate by BGS traps of *Ae. albopictus* mosquitoes in PAN and PDL at specific time periods during the baseline collection, sterile releases and monitoring activities of a SIT programme against the species in the villages from 09 February 2017 to mid-April 2018.

| Timeline | Ovitrap productivity<br>(eggs/ovitrap/week) |  |  | Egg fertility in ovitrap<br>(%) |  |  | Female capture rate<br>(females/BGS trap/day) |  |  |
| --- | --- | --- | --- | --- | --- | --- | --- | --- | --- |
|  | (mean, 95 % CI) |  | Unpaired *t* test | (mean, 95 % CI) |  | Unpaired *t* test | (mean, 95 % CI) |  | Unpaired *t* test |
|  | PAN | PDL |  | PAN | PDL |  | PAN | PDL |  |
| **Pre- sterile release** | 111 | 92 | $t = 1.98, df = 16,$ | 92.0 | 93.7 | $t = 1.47, df = 16,$ | 22 | 6 | Not done |
| **(09 Feb 2017 to mid-April 2017)** | (94 - 127) | (82 - 101) | $P = 0.066$ | (90.1 - 93.9) | (92.5 - 95.0) | $P = 0.162$ | (12 - 33) | (4 - 8) | |
| **IVM strategy and sterile release** | 36 | 28 | $t = 0.02, df = 10,$ | 71.7 | 94.2 | $t = 2.55, df = 10,$ | 5 | 2 | $t = 0.77, df = 10,$ |
| **(mid-May 2017 to mid-July 2017)** | (0 - 72) | (13 - 101) | $P = 0.985$ | (52.1 - 91.3) | (92.1 - 96.2) | $P = 0.029$ | (1 - 8) | (2 - 3) | $P = 0.457$ |
| **Sterile release before cyclonic weather** | 30 | 86 | $t = 6.09, df = 46,$ | 64.4 | 94.5 | $t = 17.13, df = 46,$ | 4 | 12 | $t = 3.15, df = 20,$ |
| **(Aug 2017 to mid-Jan 2018)** | (26 - 35) | (66 - 105) | $P < 0.0001$ | (60.5 - 68.3) | (93.7 - 95.3) | $P < 0.0001$ | (2 - 5) | (10 - 13) | $P = 0.005$ |
| **Sterile release after cyclonic weather** | 101 | 78 | $t = 0.48, df = 6,$ | 88.5 | 94.0 | $t = 0.97, df = 6,$ | 4 | 19 | Not done |
| **(mid-Jan 2018 to 09 Feb 2018)** | (44 - 158) | (41 - 115) | $P = 0.651$ | (78.7 - 98.3) | (91.8 - 96.2) | $P = 0.369$ | (0 - 11) | (0 - 53) | |
| **Post-sterile release** | 131 | 91 | $t = 1.56, df = 14,$ | 94.8 | 95.8 | $t = 1.70, df = 16,$ | 14 | 27 | $t = 2.18, df = 10,$ |
| **(09 Feb 2018 to mid-April 2018)** | (81 - 181) | (57 - 124) | $P = 0.142$ | (94.1 - 95.3) | (94.9 - 96.8) | $P = 0.104$ | (7 - 19) | (18 - 36) | $P = 0.055$ |

### Sterile releases

Two weeks after the IVM strategy stopped being implemented in the pilot sites, that is as from August 2017, the density of *Ae. albopictus* in PDL gradually increased across the weeks reaching very high levels by the end of November 2017 which coincides with the onset of the hot summer season (with a mean of 22 eggs/ovitrap/week and 7 females/BGS trap/day in the second week of August 2017 to 170 eggs/ovitrap/week and 25 female/BGS trap/day by the end of year 2017) (Fig. 1b and d). On the other hand, *Ae. albopictus* density in PAN_Out remained at very low levels throughout 2017 (not exceeding a weekly mean of 50 eggs/ovitrap/week). It rose to high levels only as from January 2018 (with a mean of 154 eggs/ovitrap/week recorded in the first week of January 2018) after the advent of persistent torrential rainfalls in the last two weeks of December 2017 (Fig. 1b and d).

Weekly mean fertility of eggs remained relatively stable in PDL throughout the study period (Fig. 1a), ranging from 91.1 % to 98.3 %, and significantly differed from PAN only during the sterile releases (*P* < 0.05, Table 2). However, the advent of the tropical cyclone Berguitta (with a pressure of 982 hPa in its centre, gusts of 140 −170 Km/h and intense precipitations of 176 mm rainfall/h) which influenced the weather in Mauritius from 15 January to 20 January 2018^16,17^ seemed to have greatly impacted the last three weeks of the release programme, since mean weekly egg fertility and ovitrap productivity did not differ significantly between PAN and PDL during that period (*P* > 0.05, Table 2) although a weekly release of 60,000 sterile males was still maintained in PAN.

Specifically for PAN, weekly mean ovitrap productivity in the village did not significantly differ from PDL before the start of the sterile releases in mid-May 2017, during the implementation of the IVM strategy (mid-May to mid-July 2017), during sterile releases in the aftermath of the tropical cyclone Berguitta (mid-January 2018 to February 2018) and after the stop of the sterile releases (February 2018 to mid-April 2018) (*P* > 0.05, Table 2). Ovitrap productivity was however significantly lower in PAN from August 2017 to mid-January 2018 (*P* < 0.0001, Table 2), that is, when release of sterile males was the only mosquito control method implemented in the village and when climatic conditions were moderate. During that time frame, weekly mean (95 % CI) induced egg sterility was 31.8 % (27.7 – 35.9 %) in PAN and a 55.7 % (46.7 - 64.6 %) decrease in ovitrap productivity was noted in the village relative to PDL. Mean capture rate of adult *Ae. albopictus* females by BGS traps was also significantly lower than in PDL (*t* = 3.151, df = 20, *P* = 0.0050) with a daily mean (95 % CI) of 4 females/BGS trap/day (2 - 5 females/BGS trap/day) and 11 females/BGS trap/day (7 - 17 females/BGS trap/day) in PAN and PDL respectively.

### Effect of ovitrap productivity on induced sterility

The Pearson’s coefficient depicting the relationship between weekly mean ovitrap productivity and egg induced sterility in PAN indicates a significant negative correlation between both parameters as illustrated by the regression analysis in Fig. 2 (*R*^2^ = 0.280; df = 1,31; *F* = 11.49; *P =* 0.002).

**Figure 2.**
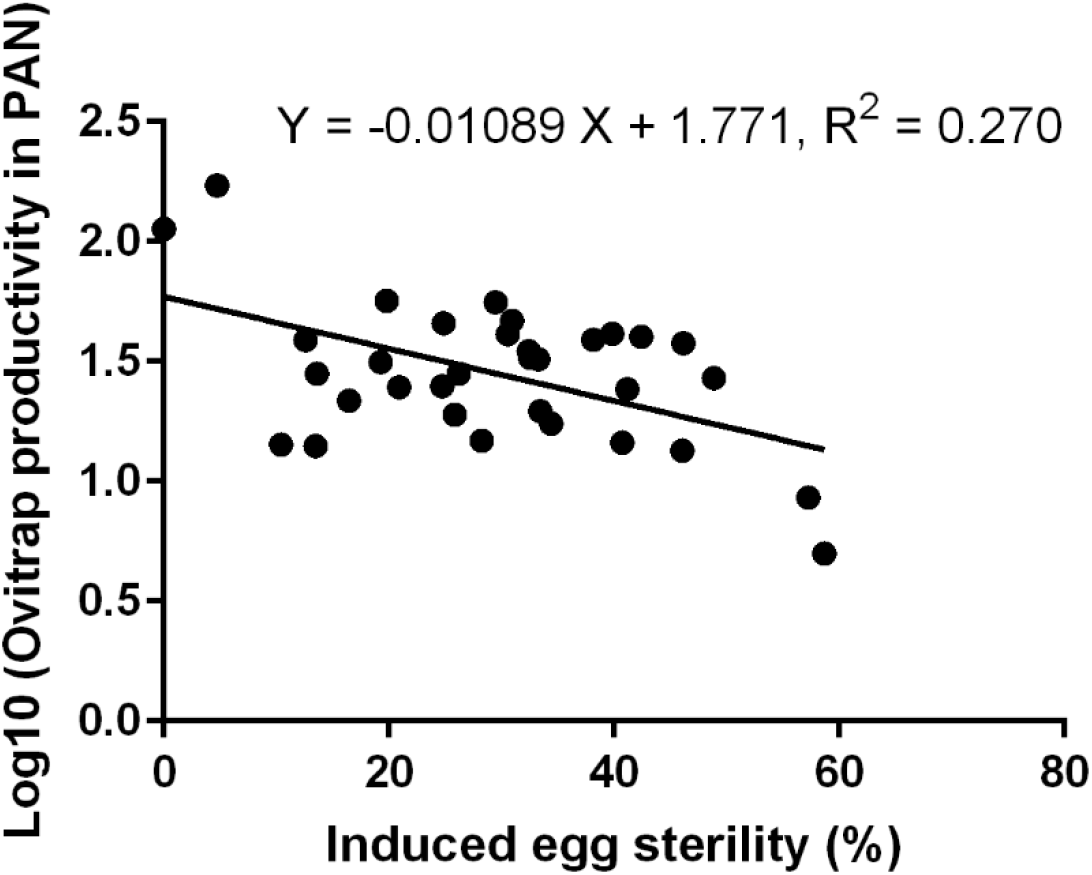
Correlation between weekly mean ovitrap productivity and induced egg sterility in PAN.

### Induced sterility and ovitrap productivity in individual ovitraps

Weekly mean induced egg sterility and ovitrap productivity of individual ovitraps calculated for the period August 2017 to mid-January 2018 respectively ranged from 0.4 to 49.4 % and from 12 to 54 eggs/ovitrap/week in PAN (including the outskirt region).

Interestingly, 50 % of ovitraps in the outskirt region, lying within a radius of 150 m from the closest release stations, had a mean induced sterility ranging from 17 to 49 %. Furthermore, ovitraps in the outskirt region tended to be more productive than those in the village and there was no clear association between induced sterility and ovitrap productivity levels within the individual ovitraps (Fig. 3).

**Figure 3.**
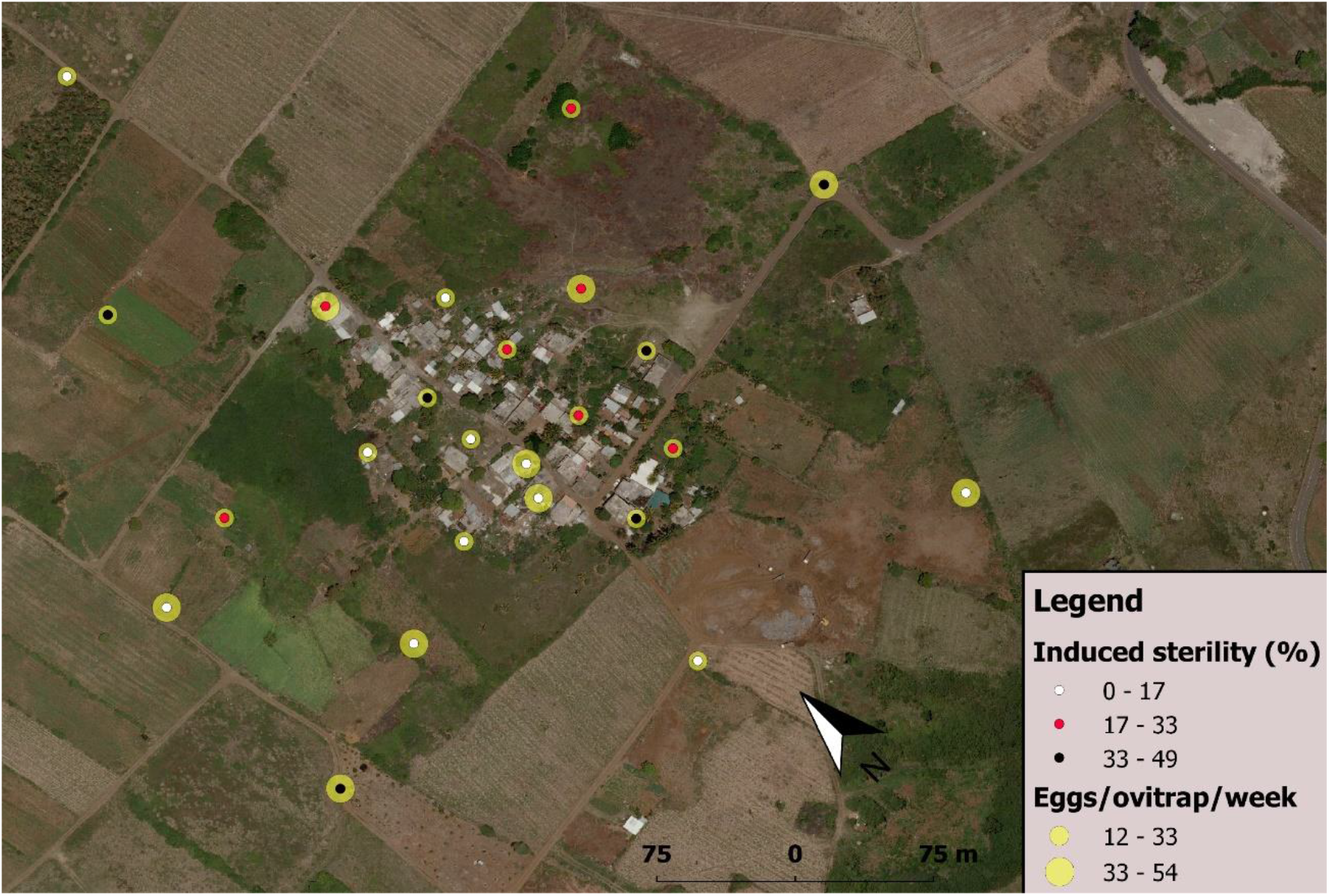
Weekly mean of induced egg sterility (%) and ovitrap productivity (eggs/ovitrap/week) in PAN during weekly release of sterile *Ae. albopictus* males from August 2017 to mid-January 2018 in the village.

### Fried’s competitiveness index

There was a strong negative correlation between the fertility rate in the release area (Ee) and the ratio of sterile to wild males (S/N) (Pearson’s product-moment correlation, cor = −0.63, p< 10^−3^). The mean (S/N) ratio during the release period was 0.53 (95 % CI 0.00-0.95) for a fertility rate of eggs of 0.81 (95 % CI 0.57-0.95) in the release area whereas the fertility rate in the control area was 0.94 (95 % CI 0.91-0.98). This corresponded to a very low recapture rate of 0.12% (SD 0.05%).

The corresponding Fried index was estimated at 0.41 (95 % CI 0.08-0.71). The point estimations of Fried index are presented against the date in Fig. 4 and no particular trend was observed (3 values out of the 23 weeks of monitoring were discarded as outliers because Ee was greater than Ha).

**Figure 4.**
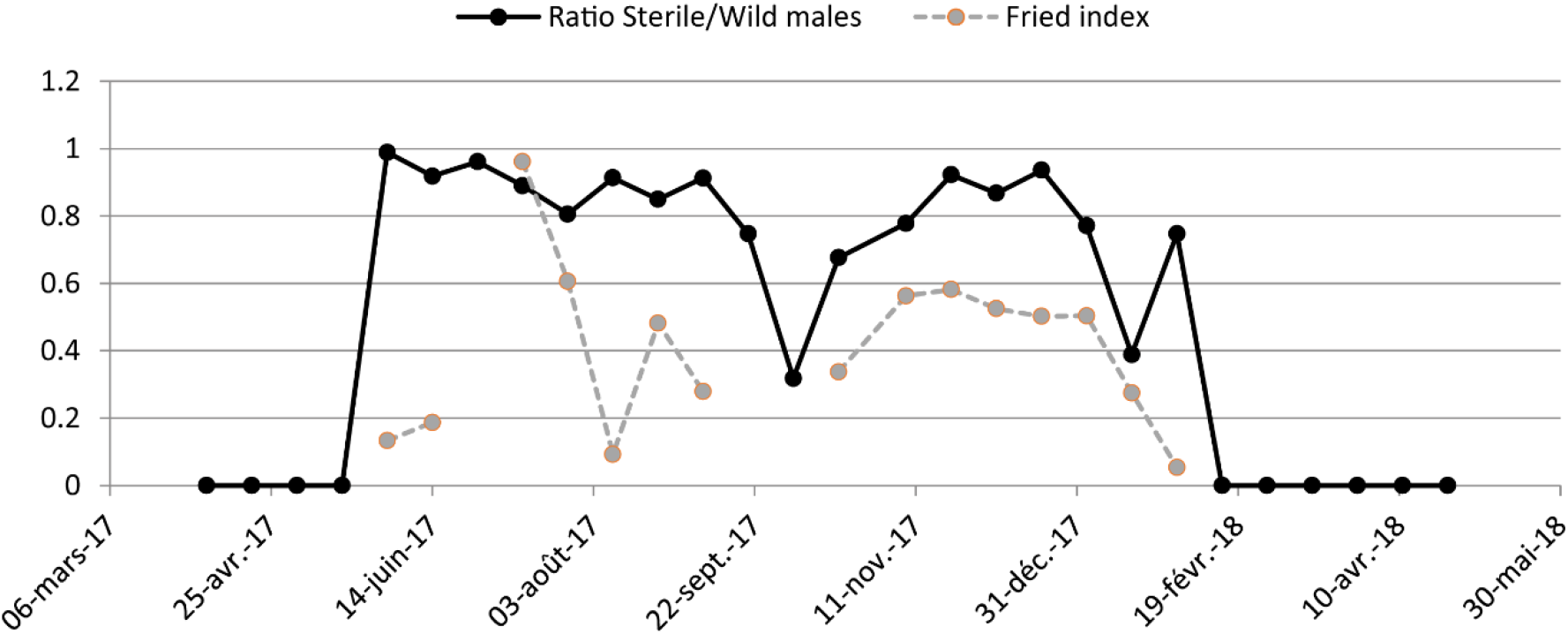
Temporal dynamics of the sterile to wild males ratio and the Fried competitiveness index in PAN during weekly release of sterile *Ae. albopictus* males from mid-May 2017 to February 2018.

### Collection of other mosquito species by BGS traps

Besides *Ae. albopictus*, other mosquito species collected by the BGS traps in the two villages include *Culex quinquefasciatus, Anopheles arabiensis, Anopheles maculipalpis* and *Aedes fowleri*. Of these, *Culex quinquefasciatus* was the most abundant species with a total of 706 and 1026 adults collected in PAN and PDL during the sterile releases. Less than 35 adults of each of the other species were collected in the two localities for the same period. Moreover, the daily mean number of adult *Culex quinquefasciatus* captured by BGS traps during the sterile releases did not differ significantly between PAN and PDL (*t* = 0.512, df = 36, *P* = 0.612).

### Cost estimation of the SIT programme

For this pilot trial, the production of 1000 sterile males costed 19 euros while the cost associated with weekly sterile male release (60 000 males over 3 ha in PAN) and monitoring (over 33 ha cumulatively for PAN and PDL) was 195 euros. In total, 582 euros were spent on a weekly basis for every hectare treated (Table 3).

**Table 3.** Cost estimation of the pilot SIT programme for *Ae. albopictus* control in PAN (including release in PAN and monitoring in PAN and PDL)

| Cost break-down of SIT trial | Euros |
| --- | --- |
| Production cost / 1000 sterile males (including quality control in laboratory and field) | 19 |
| Release and monitoring cost / ha / week | 6 |
| Number of sterile males released / ha / week | 20,000 |
| Total cost / ha / week | 582 |

## Discussion

To the knowledge of the authors, this is the second SIT programme during which radio-sterilized *Ae. albopictus* males were released to control populations of the species in the wild after that conducted in northern Italy^18^, and the first in a tropical climate. The implementation of an IVM strategy coupled with the onset of the unfavorable winter season during the first two months of the sterile release programme (mid-May to mid-July 2017) significantly reduced the density of *Ae. albopictus* in PAN and PDL. Specifically for PAN, mean weekly ovitrap productivity and mean daily capture rate of *Ae. albopictus* females from BGS traps decreased three-fold and four-fold respectively. After the cessation of the IVM strategy and during the major part of the release programme (i.e., from August 2017 to mid-January 2018), weekly release of 60,000 sterile males in PAN (corresponding to 20,000 sterile males/ha), successfully maintained the suppression of the wild *Ae. albopictus* population in the village. Mean weekly ovitrap productivity in PAN was reduced to 30 (26 - 35) eggs/ovitrap/week when sterile males were released from August 2017 to mid-January 2018 but subsequently increased by three and four folds respectively during the last three weeks of the releases in the aftermath of a cyclone and two months after the cessation of the sterile releases. It demonstrated that within specific eco-climatic conditions, SIT could be used as a control tool against *Ae. albopictus* and can, unlike conventional chemical control methods, be highly effective when the vector is found at very low densities^19^.

The discrepancy between the predicted and observed sterile/wild male release ratio (10:1 versus 1:1) in this study may have been caused by a combination of the following factors: density-dependent mortality causing an increased mortality during releases during the suppression trial in comparison to the MRR trials when lower amounts were released; increased density of the target population during the suppression trial in comparison to the density estimated during the MRR which were conducted more than two years before this suppression trial; and inaccuracy of the method used in our study to predict this ratio in the absence of marking of the sterile males. Moreover, previous MRR conducted in this area showed that the sterile males had a median dispersal distance of ~100m^20^. It is thus likely that a large part of the released sterile males dispersed to an additional area of more than 10 ha around the 3ha where the males were actually monitored. Given the small non isolated release area, the dose of sterile males per ha was thus actually between 4,600 and 20,000 males/ha/week. This could partially explain the low observed ratio, given that the densities of wild males were estimated to 3344 (95% CI, 2258–4429) ^20^. This would also explain partially the low recapture rate in the monitored area as well as the intermediary induced sterility measured in the buffer area. More studies will be needed to determinate what were the most important factors driving the low observed ratio. During a similar SIT programme releasing radio-sterilized *Ae. albopictus* males in five urbanized localities in Italy, there were no significant reduction in the wild population density in three of the four localities^18^ although the ratio of sterile to fertile male releases were high. In Boschi (16 ha), the fourth locality, mean seasonal sterile/fertile male release ratio was 350:1 and 130:1 in 2008 and 2009 respectively and percentage reduction in ovitrap density relative to the control site were 50.7 and 72.4 %. As speculated by Bellini *et al*.^18^, density of the wild *Ae. albopictus* population at the start of the releases in Italy, may have been an important determining factor in the success of the releases. Ideally, the density of mosquito populations must be reduced to low levels in the target site (for instance through the implementation of an IVM strategy and/or by utilising naturally occurring seasonality) in order to maximize the success of a sterile release programme. Furthermore, while sterile males were released in the pupal stage in Italy, results from three competitiveness studies on *Ae. albopictus* demonstrated that radio-sterilized 1-day-old adult males were less competitive than older ones^21,22,23^. To maximize male performance in the field during an SIT programme, releasing sterile males in the pupal stage or as newly-emerged adults is therefore not recommended.

Mean (95% CI) induced sterility and Fried competitiveness Index during the releases in PAN were 31.8 % (27.7 – 35.9 %) and 0.40 (0.08 - 0.71) respectively which are comparatively lower than that obtained during a previous semi-field competitiveness study using an *Ae. albopictus* strain originating from PDL and PAN^23^. More specifically, at a 5:1 sterile/wild male release ratio, mean (± s.d.) induced sterility and male competitiveness index in the cages were 76.9 ± 3.3 % and 0.91 ± 0.97 respectively. This result is in line with previous studies where radio-sterilized *Ae. albopictus* males^18,24^ and transgenic sterile *Ae. aegypti* males^25,26^ performed better in semi-field condition than in the field^27^. In this study, the discrepancy in male performance between field and semi-field conditions could be attributed to several factors including variations in eco-climatic parameters as well as potential differences in morphological size^28,29,30^and inherent traits^31^ between lab-reared and wild *Ae. albopictus* mosquitoes. Overall, a field competitiveness of 0.40 can still be considered as a good result in comparison to other published trials based on transgenic mosquitoes^32^. It was however slightly lower than the 0.5-0.7 values obtained recently in China in a successful suppression effort of *Ae. albopictus* in two isolated islands, where the Sterile Insect Technique was combined to the Incompatible Insect Technique in order to reduce the amount of radiation necessary to obtain full sterility^33^. This competitiveness of our sterile males corresponded to a level of residual fertility (~3%) which is higher than recommendations of the World Health Organization and the International Atomic Energy Agency^34^. Reducing this residual fertility by increasing the dose in future operational programmes might reduce the competitiveness. It must also be acknowledged that our estimation of the competitiveness is based on a method that is less accurate than the marking of the sterile males, even if it used routinely in fruit-fly SIT programmes (Pereira R.C., pers. com.).

Immigration of sterile females from outside of PAN could also have diluted the success of the releases. Actually, 50 % of ovitraps in the outskirt region exhibited a mean induced sterility ranging between 17 and 49 % which shows that the sterile females mated within PAN migrated to the outskirt region. This also leads us to infer that immigration of fertile females into PAN could have occurred in a symmetric way, thus reducing the measured induced sterility within PAN. It is however worth mentioning that although the migration of fertile *Ae. albopictus* females from PAN_Out into the village was not substantial enough to compromise the success of the release programme (since a significant positive correlation between egg induced sterility and ovitrap productivity was observed). It may still have reduced the impact of the latter on the target population and lead to an underestimation of the competitiveness by inflating the fertility rates measured in the ovitraps.

While the density of *Ae. albopictus* was very low in PAN_Out throughout the major part of the release programme, the population however flared up and remained at very high levels with the advent of persistent torrential rains in the last week of December 2017 (i.e., approximately one month before the end of the releases). More specifically, during the last three weeks of the releases, a notable increase in egg fertility and ovitrap productivity was observed in PAN and weekly mean of both parameters did not differ significantly between PAN and PDL. It is highly probable that the tropical cyclone Berguitta which influenced the weather in Mauritius from 15 January to 20 January 2018^16,17^, could have greatly impacted the success of the sterile releases by creating numerous larval breeding sites. Under moderate weather conditions, it is known that mosquitoes can reduce the impact of raindrops because of their strong exoskeleton and low body mass^35^ and that in many flying insects, wind has a significant impact on their flight ability as well as on their emission and responsiveness to sex pheromones^36,37,38,39,40,41,42,43,44^. However, high-intensity rainfalls and strong wind velocity may decrease the life expectancy of small delicate insects^45,46,47^ such as mosquitoes, especially sterile males reared in artificial conditions. Some insects have also been found to modify their flight, courtship, mating and foraging behaviours in response to changes in barometric pressure^48,49,50^ which usually occurs under cyclonic conditions. The high density of *Ae. albopictus* in PAN_Out coupled with cyclonic winds may also have enhanced the migration of fertile females into the village. While the maximum distance travelled by *Ae. albopictus* females in some dispersal studies ranged between 150 and 800 m^51,52,53,54^, wind-mediated migration of some mosquito species has also been expounded by others^38,55,56,57,58,59,60,61,62^.

Finally, while the suppression of a mosquito species could potentially lead to an increase in the incidence of other species sharing the same ecological niche^63^, mosquito larval surveys conducted between 2007 and 2013^64^ demonstrated that *Ae. albopictus, Culex quinquefasciatus* and *Anopheles arabiensis* were the most abundant species in PAN and PDL and that the breeding sites of the three mosquitoes rarely overlapped. Moreover, the capture rate of *Cx quinquefasciatus* by BGS traps in PAN did not differ significantly from PDL during the sterile releases while only few specimens of *An. arabiensis* were collected by the traps during the study.

Estimated cost for this SIT trial programme (582 euros/ha/week) is higher than that of the pilot trial in China (54-216 euros/ha/week)^33^. The difference is probably due to the scale of our trial, which was done in a much smaller area.

## Materials and methods

Building on the experience and lessons learnt from previous mosquito SIT programmes^18,33,65^, major pre-requisites were identified and addressed to ensure that competitive radio-sterilized males were released and that an evidence-based sterile release strategy was elaborated for an optimal coverage of sterile males in PAN.

PAN (20°04’60”S 57°41’30”E) and PDL (20°05’01”S 57°42’14”E), respectively selected as release and control sites in this study, are two rural villages in the northeast of Mauritius approximately 1.6 km apart from each other. With 268 inhabitants, 67 houses and slightly inland, PAN (0.03 km^2^) is ten times smaller than PDL (0.3 km^2^) which is a coastal village of 800 inhabitants and 203 houses. Due to their close proximity, the two villages share several similar characteristics including – they both lie in the relatively flat Northern Plains of Mauritius, have similar soil strata with high water-retentive capacity, are bordered by extensive sugar cane fields with houses very close to vegetated lands, are at essentially the same elevation (15.4 ± 4.3 m and 26.5 ± 7.2 m above sea level for PDL and PAN, respectively) and share relatively similar climatic conditions although PAN is less windy than PDL, being less influenced by sea breezes. Mean summer and winter temperatures in the two villages are respectively 28°C and 22°C while rainfall averages 1200–1800 mm annually^64,66^. Moreover, results of studies investigating *Ae. albopictus* oviposition activities in PAN and PDL indicate that the seasonal population dynamics of the species in the two villages are relatively similar^64,67^.

Sterile releases were initiated in PAN at the beginning of the winter season (mid-May 2017) when the vector density was relatively low and also because the density of an *Ae. albopictus* population living in the outskirt of PAN was minimal at that time^64,67^.

Furthermore, due to an important wild population of *Ae. albopictus* estimated in PAN^20^ and PDL^68^ during Mark-Release-Recapture (MRR) studies, an IVM strategy was implemented in both villages during the first two months of the releases (i.e. from mid-May 2017 to mid-July 2017) to reduce the vector density and hence maximize the success of the sterile releases. This study thus aimed at evaluating the incremental impact of SIT as a component of an IVM strategy. We demonstrated previously that these two villages were appropriate as paired sites^67^.

Moreover, in those studies, a low dispersal of radio-sterilized males was observed while the average life expectancy of the released males ranged between 6 and 9 days^20,68^. Hence in this study, sterile males were released on a weekly basis with distance between two adjacent release stations not exceeding 80 m. Sterile males were released as 3-days-old adults since they tended to be more competitive than 1-day-old and 5-day-old adults in semi-field enclosures^21,23^. Finally, based on the estimation of the wild male population during MRR studies in PAN and the level of induced sterility in semi-field competitiveness cages^20,23^; a weekly release of 20,000 sterile males per hectare was implemented in PAN with an aim of achieving a sterile/wild male release ratio of at least 10:1.

### Integrated Vector Management (IVM) strategy

During the first two months of the sterile release programme (15 May to 15 July 2017), an IVM strategy was implemented in PAN, its outskirt region and PDL. This involved larval source reduction activities, weekly larviciding of mosquito breeding sites using *Bacillus thuringiensis israelensis* (*Bti*) and bi-weekly fogging operations with a pyrethroid-based insecticide (Aqua K-Othrine^®^, Bayer, Germany) using hand-held pulse jet thermal foggers (Igeba TF-35, Germany) by insecticide operators of the Ministry of Health and Quality of Life. Weekly release of sterile males in PAN was also initiated as from 17 May 2017.

### Sterile male production

Generations F11-F21 of a colony recently established in the laboratory of the Vector Biology and Control Division (VBCD, Ministry of Health Quality of Life, Curepipe, Mauritius) from field-collected eggs in PAN and PDL, were used for this study. Adult mosquitoes were reared at a density of 15,000 adults in large colony cages (60 x 60 x 60 cm, Bioquip, Rancho Dominguez, Ca) with constant access to a 10% sucrose solution. Each cage was blood-fed daily for 30 minutes over a period of one month by placing one large aluminium blood plate of the Hemotek system (20 cm in diameter, containing 20 ml of blood warmed at 38 °C and covered with collagen membrane) on top of the latter. Four days after the first blood feeding, 2 ovicups (10 cm in diameter) were placed inside each cage and oviposition papers replaced every two days for four weeks. Egg papers were stored in a climate-controlled room (25 ± 1.5°C, 80 ± 5% RH) within a hermetic glass chamber kept in darkness with a saturated solution of K2SO4. Eggs stored in this way could be used for up to 10 weeks.

Eggs were quantified based on their relative weight using the protocol outlined by Zheng *et al*. (2015)^69^. Hatching procedure consisted in shaking the desired amount of eggs in 170 ml warm dechlorinated tap water (28°C) for approximately 15 seconds in a 200-ml plastic container prior to their transfer in trays containing a hatching solution consisting of 0.4 g of a food-mix (made up of 28 % tuna meal, 36 % bovine liver powder and 36 % brewer’s yeast) in 1 litre dechlorinated tap water at 28°C^70^. 13,000 newly hatched larvae were subsequently estimated volumetrically and added to each of 20 trays of three IAEA mosquito rearing units previously filled with 6 liter of dechlorinated tap water^71^. A 7.5% wt:vol slurry of a diet consisting of 80% Aquatro Tilapia Pre Grower™ (Livestock Feed Limited, Mauritius) and 20% Brewer’s yeast (MP Biomedicals, LLC) was prepared and used to feed larvae^70^ in the IAEA units during the first four days of their aquatic stage such that each larva received 0.2, 0.4, 0.6 and 0.8 mg of the diet on Day 1, 2, 3 and 4. Twenty four hours after the appearance of the first pupae, the IAEA units were tilted to collect the larval-pupal mixture using a large-bottom sieve made of loosely-knitted fabric that was specifically conceived during this study (Fig. 5).

**Figure 5.**
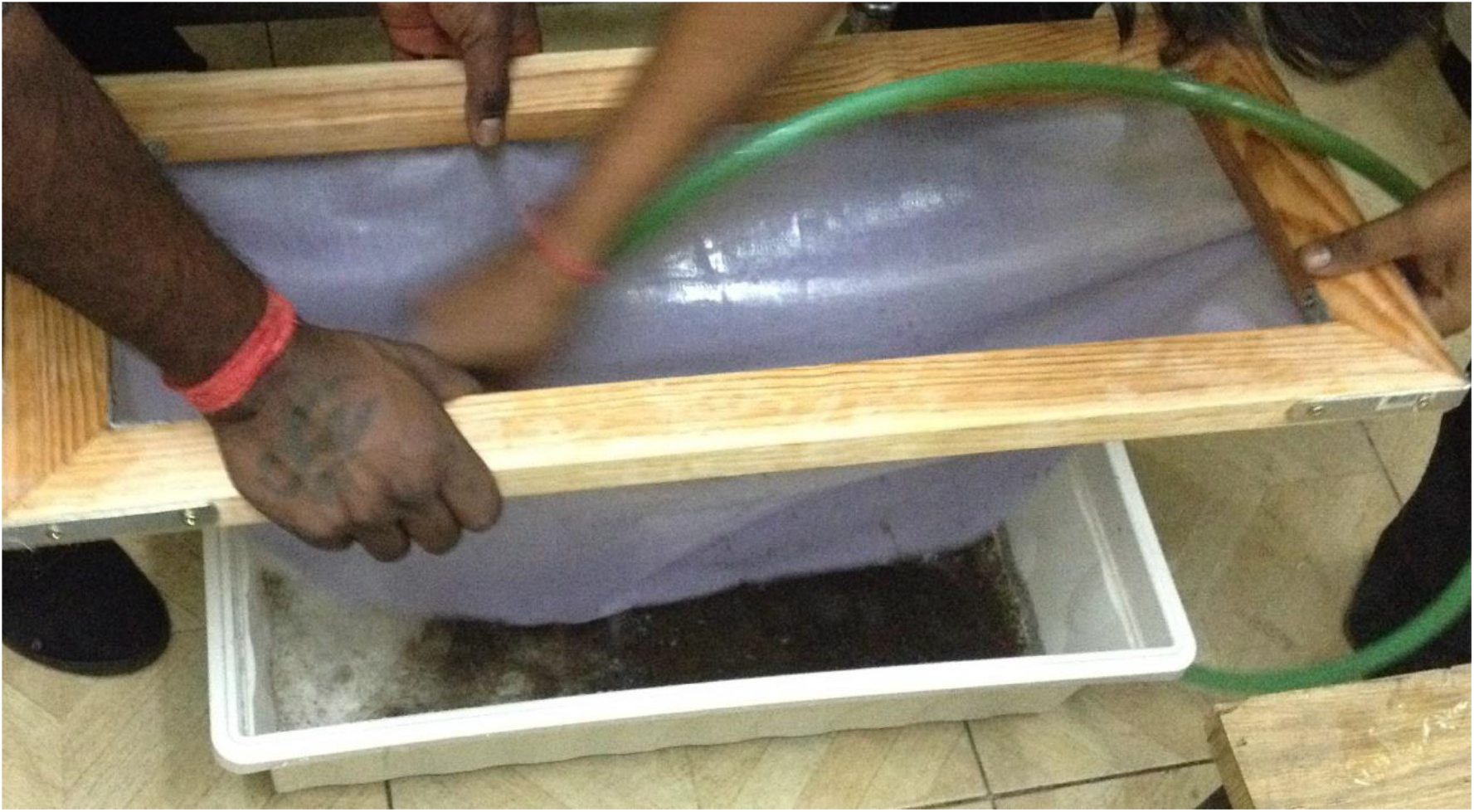
Sieve made of loosely-knitted fabric used to collect pupae from IAEA mass-rearing racks in this study.

The larval-pupal mixture collected from each IAEA unit, was equally divided into six portions and sieved separately using sieves of two different aperture sizes (1.25 and 1.4 mm). A cylinder containing the larval-pupal mixture was submerged for seven minutes in a basin of tap water heated at 37°C with two sieves of decreasing aperture (1.4 and 1.25 mm, VWR, Vienna, Austria) on top of the former. This enabled smaller (mainly male) pupae to swim to the top via the sieve apertures while larger pupae (mainly females), unable to pass through the apertures, were trapped at the bottom of the basin. To maximize the harvesting of male pupae, a second sieving step was added just after the first sieving. This consisted in passing pupae collected in the 1.4 layer through two 1.25 mm sieves for 3 minutes in a basin of tap water heated at 37°C. Pupae passing through the upper 1.25 layer were checked under a stereomicroscope and found to be mostly males. These were pooled with pupae that were collected from the 1.25 layer of the first sieving round. 15 random samples of approximately 100 pupae were taken from the pooled 1.25 layer and manually sexed under the microscope to determine the sex ratio.

Female contamination of up to 4 % were considered as acceptable in the absence of disease transmission. Male pupae thus harvested, were estimated by volume and stored in batches of 2000 per container (15 cm diameter, 8 cm height) partially filled with 50 ml water. When male pupae were 30-40 h old, they were transported to the irradiation facility in Reduit (approximately 16 Km from the VBCD) and drained of their water before irradiation.

To determine the best irradiation dose for the sterile release programme, a sterility curve was established by exposing male pupae (stored at a density of 2000 pupae per petri dish) to irradiation doses ranging from 0 to 120 Gy in a cesium-137 irradiator (Gammacell 1000, Nordion International Inc., Ontario, Canada). For each irradiation treatment, 250 irradiated males were caged with 250 virgin females for 48 h. After caging, males were removed and females blood fed daily for 30 min during 5 consecutive days. At the end of the 5 days, an ovicup was inserted in each cage and removed 2 days later. Eggs from each ovitrap paper were left to maturate for 4 days (26 ± 2°C, 70 ± 10 % RH), counted and immersed in a hatching solution (described above). The number of hatched larvae was counted 24 h after the immersion. Eggs that did not hatch were clarified with a 12 % solution of sodium hypochlorite^72,73,74^ and the number of embryos with clearly formed structures^75,76,77^ were counted under the microscope. The total number of viable eggs on an ovitrap paper was calculated as the total sum of the number of hatched larvae after the 24 h immersion in the hatching solution and the number of fully developed embryos after the clarification process. Egg fertility was calculated by expressing the total number of viable eggs as a percentage of the total number of eggs that were present on the ovitrap paper. The experiment was replicated five times. Complete sterility was achieved at 120 Gy (Fig. 6). However, based on previous *Ae. albopictus* studies reporting high competitiveness of males at 35-40 Gy^23,78,79,80,81,82^ and a marked reduction in the latter’s fitness when irradiation dose exceeded 30-40 Gy^78,80^; males were subsequently sterilized at 40 Gy for field release in this study. Mean egg fertility (95 %CI) was 3.05 (2.29 – 3.92) % in cages containing males sterilized at 40 Gy and virgin females (Fig. 6).

**Figure 6.**
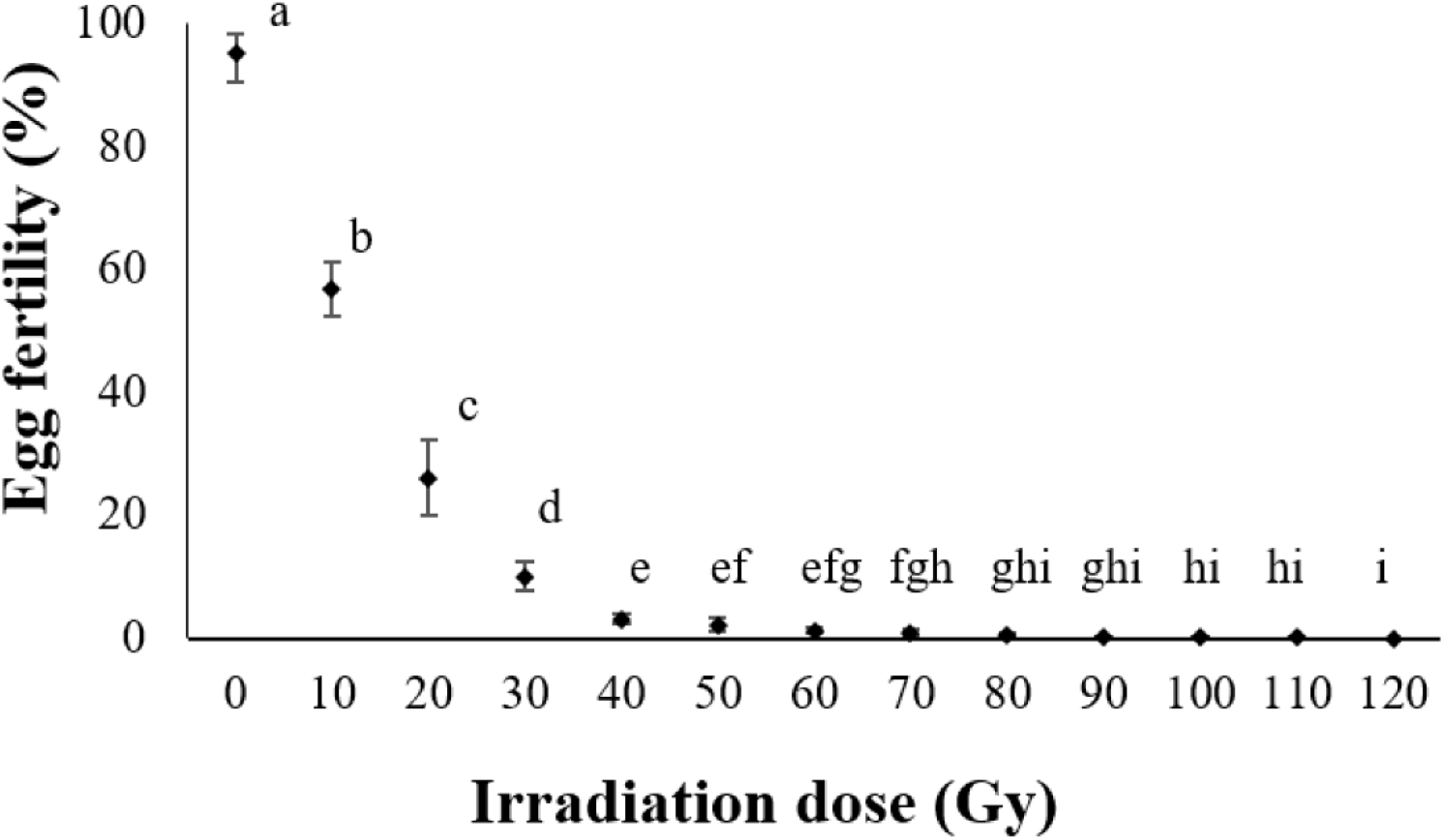
Sterility curve of lab-bred *Ae. albopictus* males. Mean egg fertility (± 95 % CI) as a function of irradiation dose. Different letters represent statistical differences between treatments (*Post hoc* Tukey tests, *P* < 0.05).

Once sterilized at 40 Gy, male pupae were transported back to the VBCD and put in Bugdorm cages (30 x 30 x 30 cm) at densities of 2000 males per cage^82^. Each cage was provided with a 10 % sucrose solution and kept in dim light at 21 ± 2°C and 70 ± 10% RH. Female contamination in the cages was removed by suction tubes until it was inferior to 1 %. When sterile males were 3-day-old adults, cages were covered with wet towels and transported in a van at ambient temperature to PAN.

### Sterile releases

On a weekly basis, 50,000 and 60,000 sterile males were released in PAN from 17 May to 31 May 2017 and from 07 June 2017 to 09 February 2018 respectively. The total number of males released was equally distributed among the release stations. Release occurred at 10 and 20 pre-determined locations in PAN (Fig. 7) from 17 May 2017 to 25 October 2017 and from 01 November 2017 to 09 February 2018 respectively. Climatic data for the region were obtained from a weather station (model Davis Vantage Pro2, Davis Instruments, USA) situated in PAN.

**Figure 7.**
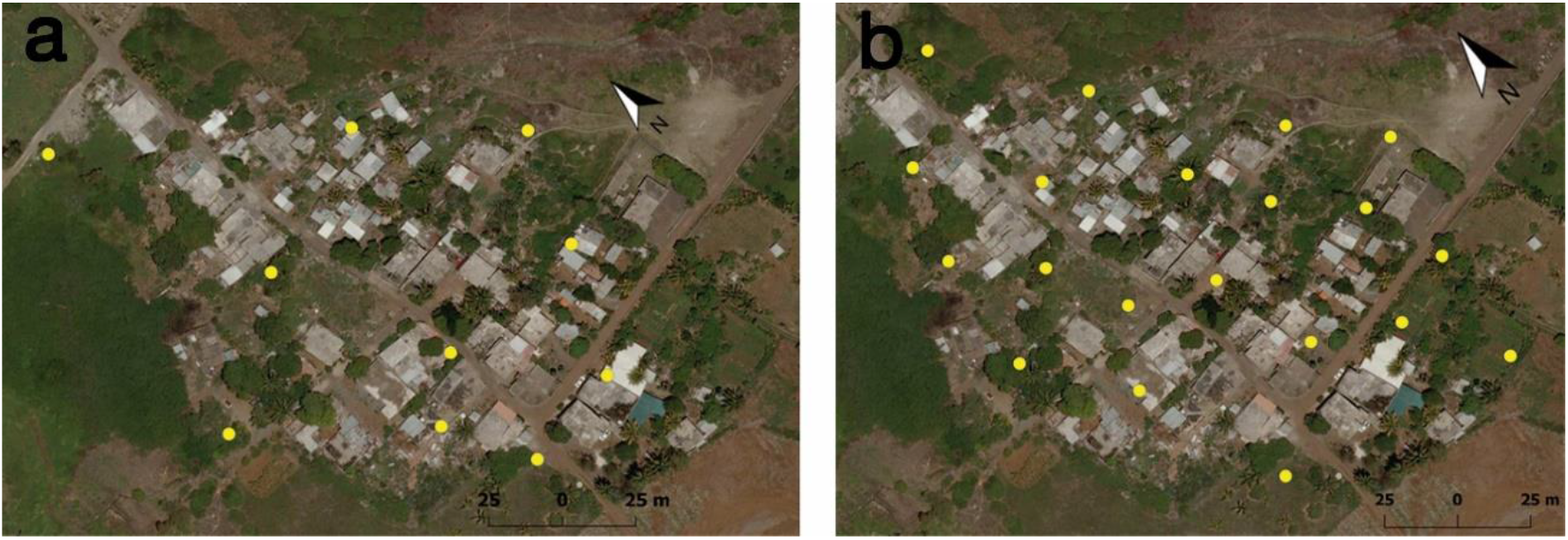
Location release stations of sterile males in PAN, **(a)** from 17 May 2017 to 25 October 2017 and **(b)** from 01 November 2017 to 09 February 2018

### Ovitrap surveillance

Before the start of the study, ovitraps were set in the villages at a large density (positioned 50 m apart) which were monitored on a weekly basis. Unproductive ovitraps or ovitraps which could not regularly be accessed were gradually removed while still ensuring a manageable yet adequate ovitrap coverage (at least 1.5 ovitraps per hectare) in the study sites. Consequently, in the present study, from 09 February 2017 to 13 April 2018, 14, 10 and 43 ovitraps were respectively placed in PAN, PAN_Out and PDL (Fig. 8). An ovitrap consisted of a black cylindrical plastic container (15 cm in height and 12 cm in diameter) filled with 400 ml tap water and lined on the inside with a strip of rough absorbent paper (34 cm x 9 cm, 145 g/m^2^, Sartorium Stedium Biotech, Goettingen) serving as an oviposition substrate. On a weekly basis, papers from the ovitraps were collected and replaced with new ones. The collected ovitrap papers were brought back to the laboratory and the number of eggs on each was counted to calculate the mean number of eggs per ovitrap per week (i.e., ovitrap productivity).

**Figure 8.**
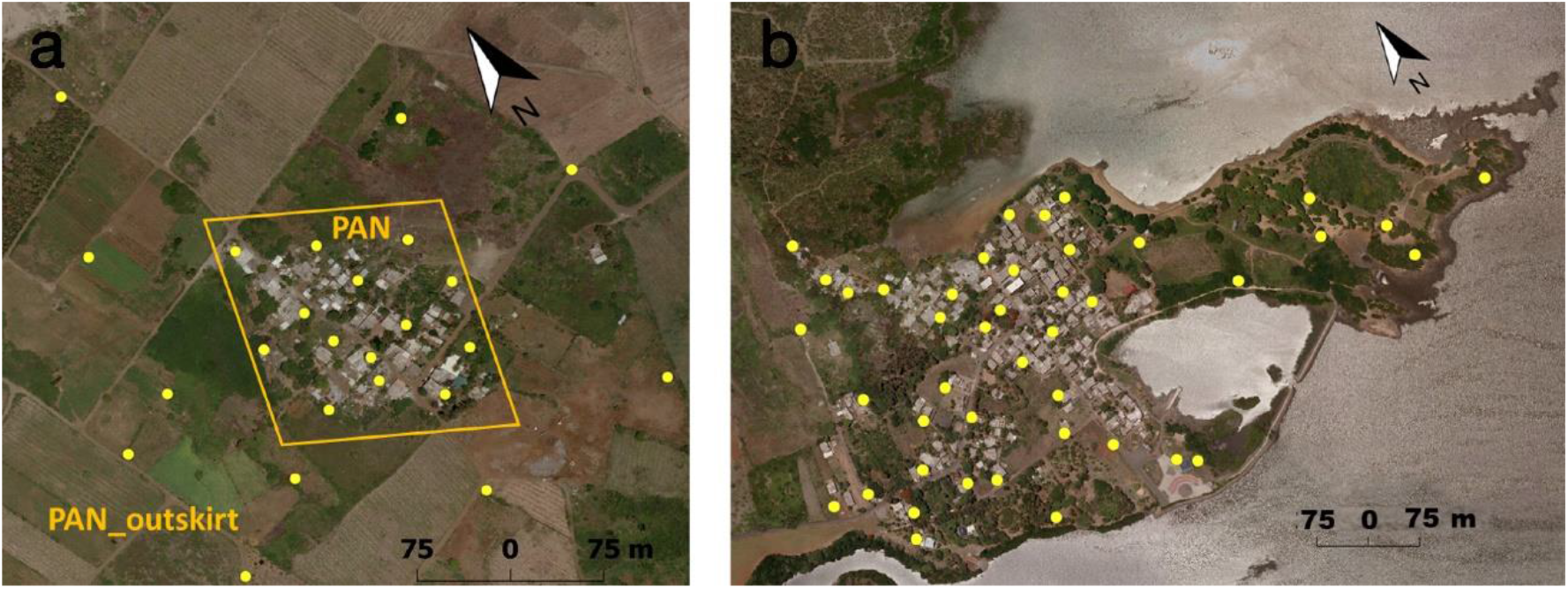
Location of ovitraps in **(A)** PAN, PAN_Out and **(B)** PDL

The majority of the eggs collected were reared to the adult stage for alimenting laboratory colonies of *Ae. albopictus* and a fraction of those adults were microscopically examined and identified as *Ae. albopictus* following MacGregor (1927)^83^ mosquito identification keys. Moreover, based on results of preliminary ovitrap surveys carried out in the study sites in 2012 during which eggs were reared to the larval stage and identified to the species level^62^, it was considered unlikely that eggs on the ovitraps papers belonged to a mosquito species other than *Ae. albopictus*.

The percentage fertility of eggs collected from all ovitraps in PAN and PAN_Out as well as from 10 of the most productive ovitraps in PDL was assessed. This consisted in counting the number of hatched and unhatched eggs on each ovitrap paper immediately after field collection and subsequently immersing the latter in a hatching solution (described above) after eggs on the paper had maturated for a period of 5 days. The number of hatched larvae was counted 24 h after the immersion. Eggs that did not hatch were clarified with a 12 % solution of sodium hypochlorite^72,73,74^ and the number of embryos with clearly formed structures^75,76,77^ were counted under the microscope. The total number of viable eggs on an ovitrap paper was calculated as the total sum of the number of eggs that had already hatched in the field, the number of hatched larvae after the 24 h immersion in the hatching solution and the number of fully developed embryos after the clarification process. Egg fertility was calculated by expressing the total number of viable eggs as a percentage of the total number of eggs that were present on the ovitrap paper.

### Adult mosquito surveillance

The adult density of *Ae. albopictus* was assessed in PAN and PDL from 05 April 2017 to 25 April 2018. On a bi-weekly basis, 8 completely black battery-operated BGS traps baited with BG-Lure (Biogents AG, Regensburg, Germany), were left to operate continuously for 24 h in each of the two villages (Fig. 9). Mosquitoes collected from the catch bags were subsequently identified and counted under a stereomicroscope following the identification keys for mosquito species outlined in MacGregor (1927)^83^. Traps were set between Wednesday and Thursday, i.e. at days 5 to 6 after release which was conducted on Friday. Since the mean survival of sterile males was previously estimated to 9 days^20^ this allowed us to consider that the ratio measured with our method is representative of what is imposed to the wild population on average.

**Figure 9.**
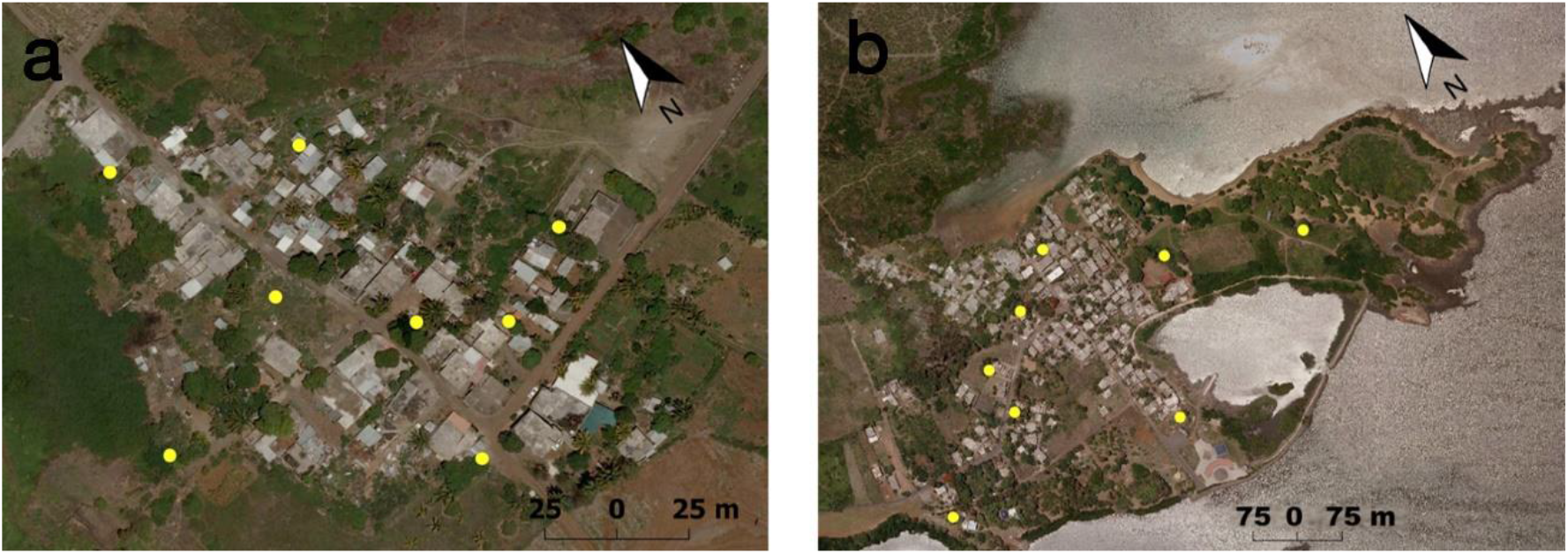
Location of BGS traps in **(A)** PAN and **(B)** PDL

### Data analysis

Weekly mean ovitrap productivity (eggs/ovitrap/week) and daily mean catches of adult female *Ae. albopictus* from BGS traps were used to estimate the temporal incidence of the mosquito in the study sites during the study. Data were tested for normality and for homogeneity of variances using respectively the Anderson-Darling and the Levene’s tests. If data were non-parametric, the latter were either angle transformed (arcsine sqrt) for frequency data or log transformed (Log10 (n+1)) for count data and tested again for normality and for homogeneity of variances prior to statistical analysis. For the sterility curve, egg fertility data were compared between treatment using analysis of variance (ANOVA) and Tukey’s post hoc tests. Spearman’s correlation coefficient (rs) was used to determine the association between weekly mean ovitrap productivity and percentage egg fertility with air temperature, and cumulative rainfall in PDL, PAN and PAN_Out. Student *t* test was used to assess for differences in weekly mean ovitrap productivity, egg fertility and daily mean *Ae. albopictus* females capture rate by BGS traps between PDL and PAN. To assess the effect of sterile males in inducing sterility in the wild *Ae. albopictus* population in PAN, the induced egg sterility value was calculated as 100 % minus the residual fertility value, which was calculated from Ho /Hn, where Ho was mean egg fertility from ovitraps in PAN during the sterile release period and Hn was the corresponding mean egg fertility from ovitraps in PDL^84,85^. The percentage decrease in ovitrap productivity (D) in PAN was calculated as follows: D = [(E_SIT_ - E_Control_) / E_Control_]^15^, where E_SIT_ and E_Control_ are the weekly mean ovitrap productivity in PAN and PDL respectively. Pearson’s correlation coefficient (*P*) and regression analysis were used to determine the association between weekly mean ovitrap productivity and induced egg sterility in PAN.

To assess the spatial effect of the sterile releases, the position of each ovitrap in PAN and PAN_Out was georeferenced using GPS - GARMIN eTrex-H and the mean ovitrap productivity and induced egg sterility of individual ovitraps calculated for August 2017 to mid-January 2018. Data obtained from the two above-mentioned variables were subsequently categorized into classes and mapped using a Geographic Information System Software (QGIS^®^ release 2.16.3).

All statistical analyses were performed using GraphPad Prism release 5.01 for Windows^®^ (GraphPad Software, San Diego USA) with alpha level of 0.05. To aid interpretability, raw data are presented in the tables and figures.

### Fried’s competitiveness index

The sterile males were not marked before release which prevented a direct measure of the sterile to wild male ratio. It was however possible to infer this ratio by comparing the male to female ratios between the two sites. Since the male to female ratio was not different between PAN and PDL outside the period of sterile male release (05 April to 03 May 2017 and 14 February to 25 April 2018) (Welch Two Sample t-test, *t* = −1.8487, df = 8.2433, *P* = 0.1006), the weekly sex ratio measured in PDL was used to infer the number of wild males corresponding to the number of wild females captured each week at the village level. This allowed predicting the number of sterile males captured, by deducting the number of wild males inferred in this way to the total number of males captured.

This ratio was used to implement Fried index^86^. This index compares fertility rates of eggs in wild females in the absence or presence of sterile males, taking into account the sterile to wild male ratio. The Fried competitiveness index was calculated using the following formula: 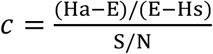, with Ha being the percentage of fertile eggs in the control area, E the percentage of fertile eggs during the same week in the release site, Hs the hatch rate from eggs of females mated with sterile males as estimated from a recent semi-field study conducted in Mauritius where egg fertility (mean ± SD) was 3.06 ± 0.17 % in control cages containing 40 Gy-sterilized males and virgin females^82^ and S/N the sterile to wild male ratio calculated as presented above.

## Data Availability Statement

All raw data are available as a Supplementary File.

## Acknowledgements

This study is sponsored by the International Atomic Energy Agency (IAEA) under MAR 5019, the Coordinated Research Programme D44002 as well as by the Joint Food and Agriculture Organization of the United Nations/International Atomic Energy Agency (FAO/IAEA) Division of Nuclear Techniques in Food and Agriculture. The authors are also thankful to the Ministry of Health and Quality of Life, Mauritius for providing transport and infrastructural logistic to carry out this research and to staffs of the Vector Biology and Control Division for their participation in ovitrap surveys. We thank Khouaildi Elahee, Nabiihah Munglee and Surendra Puryag for assistance in the laboratory.

## Author Contributions

D.P.I., A.B. and S.F. designed the study. A.B. and S.F. supervised the research. D.P.I. and J.B. analysed the data and wrote the manuscript. All authors reviewed the manuscript.

## Additional Information

### Supplementary information

None

### Competing Interests

The authors declare no competing interests.

## Notes

### Competing Interest Statement

The authors have declared no competing interest.

